# Penicillium hordei acidification precipitates Bacillus subtilis lipopeptides to evade inhibition

**DOI:** 10.1101/2025.05.21.655424

**Authors:** Manca Vertot, Morten D. Schostag, Aaron J. C. Andersen, Jens C. Frisvad, Carlos N. Lozano-Andrade, Scott A. Jarmusch

## Abstract

Interkingdom interactions are crucial for community and ecosystem functioning, however secondary metabolites mediating interactions between plant beneficial bacteria and fungi remain understudied. *Penicillium* and *Bacillus* species can individually suppress soilborne phytopathogens and promote plant growth. Here, we showed that *Penicillium hordei* and *Bacillus subtilis* co-culture led to precipitation of *B. subtilis* lipopeptides, observed as white line in agar. Metabolomic analysis revealed *B. subtilis* triggered enhanced production of fungal terrestric acid and its biosynthetic intermediates, which induced lipopeptide precipitation to prevent *P. hordei* inhibition by chemical inactivation and physical barrier formation. Besides lipopeptide precipitation, terrestric acid-mediated acidification progressively reduced production of antifungal plipastatins. The lack of lipopeptide production permitted *P. hordei* to invade and overgrow *B. subtilis* colony. We demonstrated that the white line phenomenon was conserved among closely related fungi via secretion of terrestric, fulvic or barceloneic acids.

Furthermore, terrestric acid at specific concentrations acted as a universal metabolite that drives *B. subtilis* lipopeptide precipitation even in distantly related fungi. This study provides new insights into acidification as a fungal defensive strategy that may promote co-existence with beneficial bacteria exhibiting strong antagonistic potential, thereby contributing to the formation of a stable rhizosphere community.

## Introduction

Fungi and bacteria are commonly found co-occurring in diverse habitats, including densely populated and nutrient-enriched layers of soil surrounding the plant roots (i.e. the rhizosphere) [1]. In the rhizosphere, they form complex microbial communities with other microorganisms and play a pivotal role in shaping community structure and function that ultimately affects plant health and ecosystem productivity [2,3]. Their co-existence leads to extensive interkingdom interactions, ranging from mutualistic cooperation to antagonistic competition, which are mediated by small diffusible molecules known as secondary metabolites (SMs) [4–6]. While soil fungi and bacteria have been extensively studied individually for their production of SMs, little is known about the role these SMs play during interactions with one another.

*Penicillium hordei* resides in *Penicillium* section *Fasciculata*, series *Corymbifera* [7], characterized by high extracellular enzyme activity, growth at low temperatures and close association with the rhizosphere of vegetables and flower bulbs. Unlike other members of this series, *P. hordei* does not cause blue mould storage rot and it is uniquely associated with the rhizosphere of barley and other cereals [8,9]. It is a prolific producer of pharmaceutically important ergot alkaloids, roquefortines and organic acids [10,11]. Organic acids have been shown to confer a competitive advantage to producing microorganisms. For example, they can enhance the activity of cell wall degrading enzymes, stimulate the accumulation of SMs required for pathogenesis and counteract *Fusarium*-induced virulence [12–14]. However, their ecological relevance in microbial interactions is often overshadowed by more prominent SMs such as antibiotics.

*Bacillus subtilis* is a soil-dwelling bacterial species known to promote plant growth and act as a biocontrol agent against a wide range of organisms, such as bacteria [15], nematodes [16], and fungi [17,18]. This capacity is attributed to its prolific biosynthetic potential to produce chemically diverse SMs, including cyclic lipopeptides (LPs), polyketides, and ribosomally synthesized and post-transcriptionally modified peptides. Among these, the LPs families surfactins and plipastatins have attracted the most attention due to their applications in biotechnological industry [19,20]. From an ecological perspective, these SMs improve *Bacillus* fitness in the rhizosphere by enabling the bacterium to effectively colonize roots, form biofilm and persist in the niche [21]. Furthermore, they also serve as defense metabolites against cohabiting competitors, mediate microbial communication and facilitate nutrient acquisition [22–24]. For example, surfactin-dependent motility allows *B. subtilis* to evade harmful metabolites produced by competitors [25,26]. Similarly, Kiesewalter et al. showed plipastatins and surfactins are required to hinder the growth of *Botrytis cinerea* [27]. Microbial interactions are considered a major factor in modulating *Bacillus* lipopeptide production [28]. While research has primarily focused on interactions with phytopathogens, the role of these SMs in interactions between *Bacillus* and epiphytic, non-pathogenic fungi remains largely unexplored.

In this study, we investigated the chemical interplay between prominent rhizosphere inhabitants *P. hordei* and *B. subtilis* . By combining liquid chromatography coupled with mass spectrometry-based metabolomics, mass spectrometry imaging (MSI) and molecular approaches, we demonstrated that *P. hordei* co-cultivation with *B. subtilis* led to enhanced production of terrestric acid and its biosynthetic derivatives. Consequently, these acids drove *Bacillus* lipopeptide precipitation (observed as a white line) to confer a protective mechanism for the fungus. In addition to lipopeptide precipitation, terrestric acid-mediated acidification reduced the production of antifungal plipastatins by *B*. *subtilis*. The loss of lipopeptide production allowed *P. hordei* to overgrow *B. subtilis,* highlighting the vital role of *Bacillus* lipopeptidome in sustaining competitive fitness to thrive upon co-culture with the fungus. We further revealed that the white line precipitation phenomenon was conserved among closely related species through the production of different organic acids, as well as in distantly related fungi capable of producing terrestric acid, pointing to a widespread distribution of such defense strategy. Overall, our data suggests the complex interplay between these two beneficial soil microbes is mediated via SMs and may play an important role in the rhizosphere.

## Materials and Methods

### Fungal and bacterial strains and culture conditions

Bacterial and fungal strains used in this study are listed in Table S1 and Table S2 in the supplemental material. *B. subtilis* strains were routinely grown in lysogeny broth (LB)-Lennox medium (10 g L ^-1^ tryptone, 5 g L ^-1^ yeast extract, and 5 g L ^-1^ NaCl) at 37 °C with shaking (180 rpm) and antibiotics when necessary. The final antibiotic concentrations for *B. subtilis* were: macrolide-lincosamide-streptogramin B (MLS) antibiotics (1lilμg mL^-1^ erythromycin and 12.5lilμg mL^-1^ lincomycin), spectinomycin (100lilμg mL^-1^) and tetracycline (10lilμg mL^-1^).

All fungal strains were routinely cultured on BD Difco potato dextrose agar (PDA) medium. After incubation for 7 days at 25 °C, spores were collected in 20% glycerol/water stock for subsequent experiments. To reduce terrestric acid production by *P. hordei* , LB supplemented with 20% maltose was used.

### Construction of *B. subtilis* P5_B1 *srfAC* Δ*ppsC*

Mutant strain was obtained using natural competence by transforming genomic DNA (gDNA) and selecting for antibiotic resistance as previously described [29]. Briefly, gDNA from a donor strain 3610 Δ*ppsC* was extracted using Monarch® Genomic DNA Purification Kit (NEB) following manufacturer’s instructions and 100 ng gDNA was added to P5_B1 *srfAC* grown in 200 μL at 37 °C in competence medium 80 mM K _2_HPO_4_, 38.2 mM KH _2_PO_4_, 20 g L ^-1^ glucose, 3 mM tri-Na citrate, 45 μM ferric ammonium citrate, 1 g L ^-1^ casein hydrolysate, 2 g L ^-1^ K-glutamate, 0.335 μM MgSO_4_·7H_2_O, 0.005% [w/v] tryptophan). Cells were incubated with gDNA at 37 °C for 4 h before 50 μL of the transformation mix was spread onto LB agar supplemented with spectinomycin and tetracycline and incubated at 37 °C for 24 h. Mutant strain was validated by LC-MS analysis.

### Pairwise interactions on agar

For co-cultivation, overnight cultures of *B. subtilis* strains were normalized to an optical density at 600 nm (OD _600_) of 1 in LB. Subsequently, 1 µL of bacterial suspension and 1 µL of *P. hordei* suspension were applied at a 2.5 cm distance onto PDA and PDA supplemented with bromocresol purple (0.02 g L ^-1^) and adjusted to pH 7. For each co-cultivation condition, a corresponding monoculture with the same inoculum was prepared. Three biological replicates of each condition were plated and incubated for 3 days, 5 days, 7 days, 9 days and 12 days at 25 °C in darkness.

The same co-cultivation process was applied for interactions with other *Penicillium* strains (Table S2) and LP-producing bacteria (Table S1). Plates were incubated for 9 days at 25 °C in darkness.

### Extraction of secondary metabolites

To study SMs produced by *B. subtilis* strain*s* and *P. hordei* and how they vary in response to co-cultivation we performed an untargeted time-course metabolomic analysis with four time points: days 3, 5, 7, and 9. For each time point, a 4 cm × 2 cm area of agar containing the co-cultures and a 2 cm × 2 cm area for the monocultures were cut out, sliced into small pieces, and transferred into 15 ml tubes. The cultures were extracted with 1:3 isopropanol:ethyl acetate acidified with 1% formic acid and dried under nitrogen flow. For LC-MS/MS analysis, the dried extracts were dissolved in 1.2 mL methanol (MeOH). The same extraction procedure was applied to interactions with other closely related *Penicillium* strains.

### LC-MS/MS data acquisition and processing

Crude extracts were analysed by an Agilent 1290 Infinity II UHPLC (Agilent Technologies; Santa Clara, CA) equipped with an Agilent Poroshell 120 Phenyl Hexyl column (1.7 µm, 150 × 2.1 mm) coupled to a Bruker TIMS TOF Flex. Chromatographic separation of extracts was achieved using the following solvent system: acetonitrile and water (both containing 20 mM formic acid), a flow rate of 0.35 mL min ^-1^, a column temperature of 40 °C and gradient over 14 min. The gradient started at 10% acetonitrile, increasing to 100% over 10 min, held at 100% for 2 min and returned to 10% acetonitrile over 0.1 min. The scan range was set to m/z 100-1400 and spectra were acquired at a rate of 8 Hz in both the MS and MS/MS level, in negative and positive mode. Automated DDA fragmentation was used with multiple collision energies (20 and 40 eV).

After LC-MS/MS acquisition, raw spectra were converted to .mzXML files using Bruker DataAnalysis and preprocessed using MZmine 3 [30]. Molecular networking and library search were performed within the GNPS platform [31] using feature-based molecular networking workflow (FBMN) [32].

### Sample preparation and matrix-assisted laser desorption ionization imaging mass spectrometry (MALDI-MSI)

Co-cultures and monocultures were prepared as described above onto a 10 mL PDA with bromocresol purple at 25 °C in darkness. After 9 days of incubation, the aerial hyphae of *P. hordei* were gently removed with a cotton swab dampened in water (H_2_O). Afterwards, a region of agar containing either *P. hordei* and a *Bacillus* strain in co-culture, or a singly grown microbial culture, was excised and placed onto a Bruker IntelliSlide coated with a layer of glue applied using a glue pen. Samples were dried for 2-5 hours at 36 °C followed by matrix application (30 mg mL ^-1^ 2,5-dihydroxybenzoic acid, 50:50:0.1% H _2_O:MeOH:trifluoroacetic acid) for 12 passes using HTX-Sprayer (HTX Imaging, Chapel Hill, NC, USA) at 70 °C. Samples were dried in a desiccator overnight prior to MSI measurement. The samples were then subjected to a Bruker TIMS TOF Flex (Bruker Daltonik GmbH, Bremen, GE) mass spectrometer for MALDI MSI acquisition in positive MS scan mode with 60 µm raster width and a *m/z* range of 100-2000. Calibration was done using red phosphorus. The settings in the timsControl were as follow: Laser: imaging 60 µm, Power Boost 1.0%, scan range 56 µm in the XY interval, and laser power 80%; Tune: Funnel 1 RF 300 Vpp, Funnel 2 RF 300 Vpp, Multipole RF 300 Vpp, isCID 0 eV, Deflection Delta 70 V, MALDI plate offset 100 V, quadrupole ion energy 5 eV, quadrupole low mass 100 *m/z*, collision energy 10 eV, focus pre TOF transfer time 75 µs, pre-pulse storage 8 µs. Data was normalized to the total ion count and processed as previously described by Lyng et al. [33].

### Growth monitoring of *Bacillus* strains supplemented with terrestric acid

Growth experiments were performed in a 96-well microplate. Each well contained 160 µl of LB medium supplemented with 20 µL of terrestric acid at five concentrations (400, 200, 100, 50, and 25 µg mL ^-1^), and 20 µL of either the *B. subtilis* wild type or its mutants were added. An untreated control containing only LB and bacterial suspension, as well as solvent control with MeOH and bacterial suspension, were included in the experiment. Four biological replicates were performed for each strain. The effect of terrestric acid was estimated by measuring the optical density at 600 nm every 5 min for 24 h with BioTek Synergy HTX Multimode Reader (Agilent), with continuous shaking at 150 rpm at 30 °C.

### Statistical analysis

Data analysis and visualization were performed using R 4.2.3 and the package ggplot2 [34]. Statistical analysis for pairwise comparison was explored via Student’s t or Welch’s test. For multiple comparisons (more than two treatments), one-way analysis of variance (ANOVA) followed by post hoc Tukey’s Honest Significant Distance (HSD) were performed. In all the cases, normality and equal variance were assessed using the Shapiro–Wilk and Levene tests. Statistical significance (α) was set at 0.05.

## Results

### Co-cultivation of *P. hordei* and *B. subtilis* leads to lipopeptide precipitation

To explore the chemical interplay driving cross-kingdom interactions between fungi and bacteria, we co-cultivated two beneficial wild rhizosphere isolates, *P. hordei* and *B. subtilis* P5_B1, on PDA. After 6 days of co-cultivation, a white precipitate formed near the *B. subtilis* colony. The precipitation progressively increased over time, expanding outward from the interface and gradually surrounding the *Bacillus* colony (Fig. 1A; Video S1). To characterize the chemical composition of the white precipitate, the agar between the colonies was excised, extracted and analyzed by LC-MS/MS. Initial profiling of crude extracts revealed a mixture of fungal organic acids and *Bacillus* cyclic lipopeptides. Mass spectrometry-based fragmentation library matching and feature-based molecular networking identified the organic acids as terrestric acid (TA) (1) and its biosynthetic byproducts: dehydroterrestric acid (deTA), crustosic acid (CA) (2) and viridicatic acid (VA) (Fig. 1C, Fig. S1). Similarly, the *B. subtilis* lipopeptides (LPs) were identified as plipastatin (plipastatin A2, B1, B2) (3) and surfactin isoforms (surfactin C13-C15) (4) (Fig. 1C, Fig. 1D), all common lipopeptides produced by several strains of *B. subtilis* [20].

**Fig. 1.**
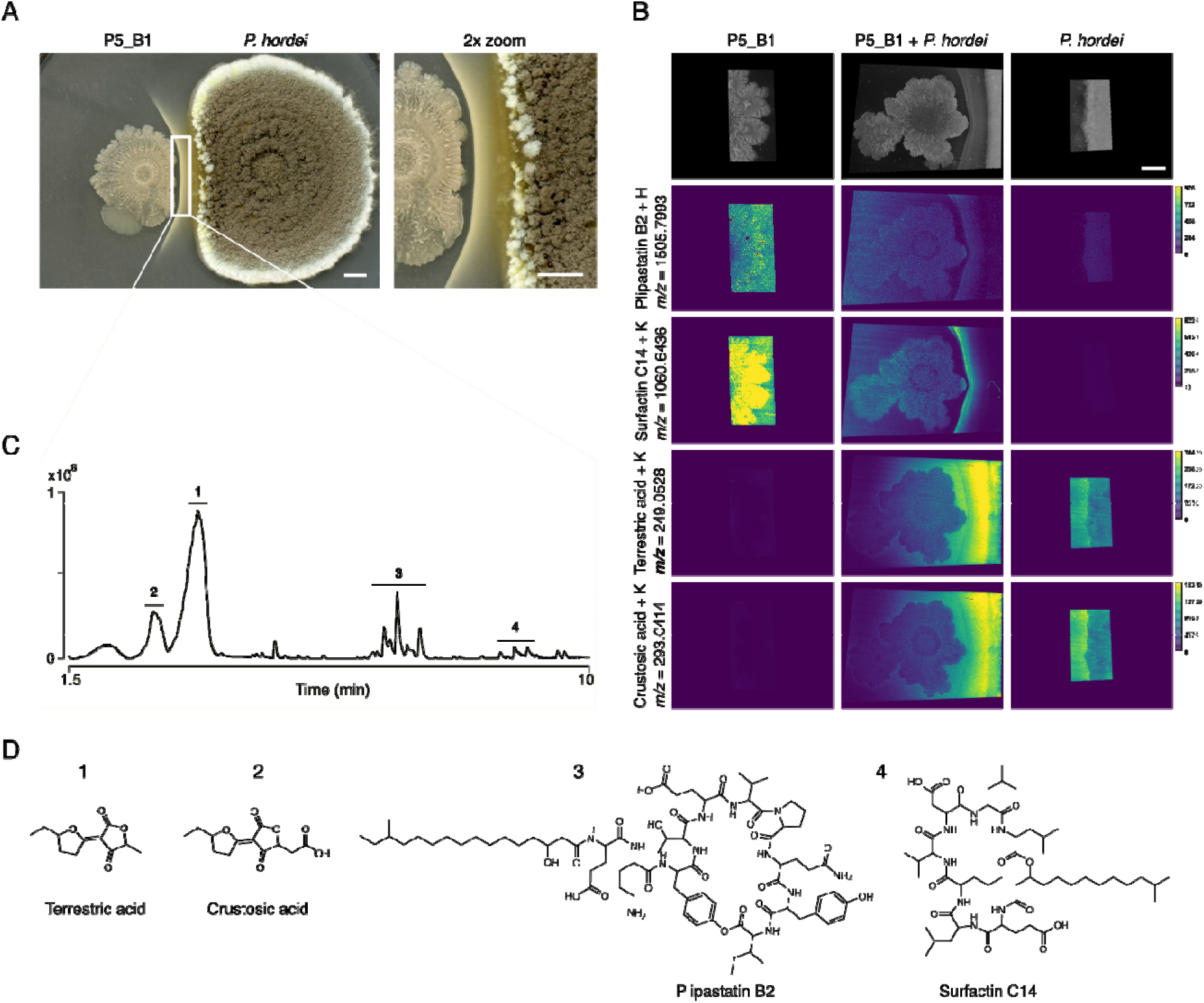
*P. hordei* induces precipitation of *B. subtilis* lipopeptides. **(A)** Interaction between *P. hordei* (right colony) and *B. subtilis* P5_B1 (left colony) on PDA plate results in white precipitate formation. Close-up (2x) of interacting area is shown on the right, and image was taken on day 12. The square indicates the area selected for LC-MS/MS analysis shown in panel C. Scale bar = 5 mm. **B** MALDI-MSI reveals the spatial distribution of selected ions in the interaction and corresponding monocultures on day 9. Ion images are shown in viridis color scheme and colored scale bars besides ion images indicate total ion count. The complete MALDI-MSI data set is shown in Fig. S2. Scale bar = 5 mm (top-right image). **C** Base peak chromatogram (BPC) in the positive mode of the crude extract from the white precipitate region area showing the accumulation of fungal organic acids and bacterial cyclic lipopeptides. The y-axis represents peak intensities. Numbers correspond to the following SMs: 1: terrestric acid; 2: crustosic acid; 3: plipastatins; 4: surfactins. D Chemical structures of SMs detected from the white line precipitate region. Surfactin C14 and plipastatin B2 are shown as isoform representatives for surfactins and plipastatins.

To resolve the spatial distribution of the SMs observed in the bulk extraction, the interacting *P. hordei* and *B. subtillis* P5_B1 colonies, along with their corresponding monocultures, were excised from thin agar plates on day 9 and subjected to MALDI -MSI. MSI confirmed the accumulation of *B. subtilis* lipopeptides (surfactins, plipastatins, and gageopeptides) in the region corresponding to the white line precipitate (Fig. 1B). Specifically, ions at *m/z* 1030.6360, 1060.6436, and 1074.6563 were identified as potassium adducts of surfactins with fatty acid chain lengths of C13, C14, and C15, respectively (Fig. 1B, Fig. S2). These surfactin isoforms exhibited the highest signal intensity at the white precipitate region and in the *B. subtilis* colony, particularly on the side facing away from the interaction zone. Similarly, isoforms belonging to the plipastatin family ( *m/z* 1491.8231 and 1505.7993) and the linear LP gageopeptides (*m/z* 702.4742, 738.4721, and 754.4456) followed the same spatial distribution pattern. However, plipastatin relative accumulation in this area was less pronounced compared to the surfactins (Fig. 1B, Fig. S2).

TA and CA were also detected, showing enhanced production upon interaction with *B. subtilis*. These ions diffused from the *P. hordei* colony across the *B. subtilis* colony, extending nearly to its far edge (Fig. 1B). The highest signal intensity was observed at the interface, spanning from the fungal colony to the white line precipitate, suggesting that the precipitate may act as a physical barrier, blocking further diffusion of TA and ultimately both fungal and bacterial growth towards each other. The presence of LPs in this region, along with the high relative concentration of TA and its biosynthetic byproducts, led us to hypothesize that organic acids secreted by *P. hordei* induce the precipitation of antifungal lipopeptides produced by *B. subtilis*, thereby conferring a protective effect on the fungus.

### Lipopeptide precipitation induced by *P. hordei* acidification prevents fungal inhibition

We visually assessed the ability of *P. hordei* to alter the pH of culture media using two pH indicators, bromocresol purple (purple when pH is above 6.8 and yellow below 5.2) and bromocresol green (blue to yellow below pH 3.8). When cultured alone on these plates, *P. hordei* triggered a rapid and progressive drop in pH on bromocresol purple within 7 days (Fig. S3A). By day 7, bromocresol green plates confirmed that *P. hordei* lowered the pH of the medium below 3.8, demonstrating a strong acidification capacity (Fig. S3B). This observation aligns with our previous LC-MS/MS analyses, which identified TA as the primary small organic acid secreted by *P. hordei* in interaction with *B. subtilis* (Fig. 1C).

Further, we explored how acid production drives the LP precipitation by combining the use of pH indicators plates with untargeted metabolomics analysis, allowing for visual inspection of LP precipitation and relative quantification of SMs in the interacting region over time (Fig. 2A).

**Fig. 2.**
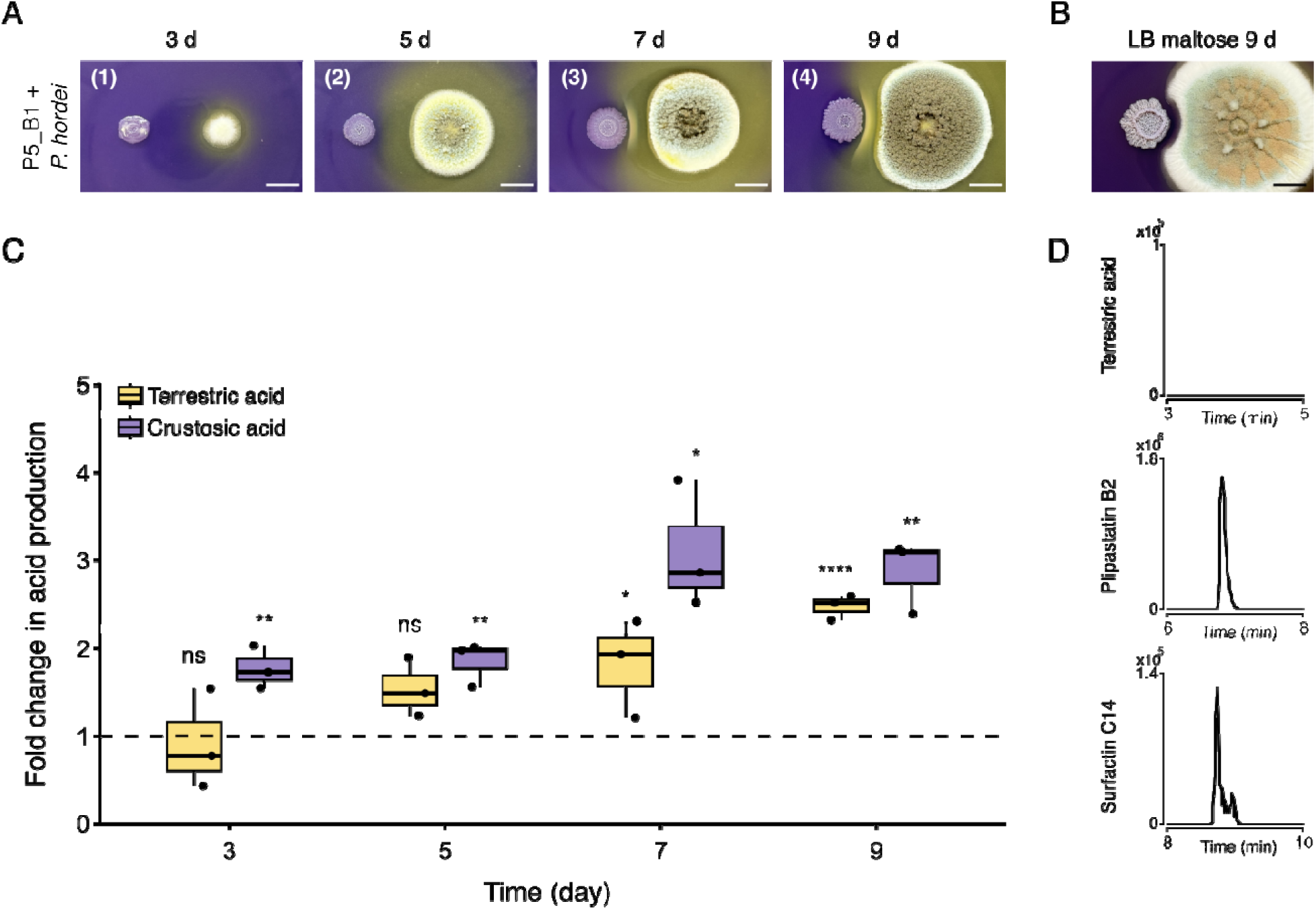
Enhanced production of organic acids by *P. hordei* drives *B. subtilis* lipopeptide precipitation to prevent fungal inhibition. **A** Time-course of the interaction between *P. hordei* and *B. subtilis* on pH indicator plates. Numbers in the upper left corner correspond to different phases of interaction: (1) before white line formation; (2) before white line formation but with reduced distance between colonies; (3) onset of lipopeptide precipitation and (4) accumulation of lipopeptides, leading to theformation of a thicker white precipitate. Scale bar = 1 cm. B *P. hordei* failed to induce lipopeptide precipitation when acid production was highly reduced in interaction on LB containing 20% maltose by day 9. Scale bar = 1 cm C Impact on terrestric (yellow) and crustosic (purple) acid production by *P. hordei* in response to *B. subtilis* over time (days 3, 5, 7, 9) on pH indicator plates. Fold changes were calculated based on relative quantification of the acids by LC-MS (peak area) in co-culture compared to monoculture of *P. hordei* (fold change = 1, dashed line). Box plots were generated from three biological replicates (n = 3), where whiskers span from the minimum to maximum values, and black line inside the box indicates median. Statistical significance for terrestric acid production was calculated using Student’s t-test, while Welch’s t-test was used for crustosic acid production, where “ns” denotes no significant difference; "*“, *P* < 0.05; "**“, *P* < 0.01; "***“, *P* < 0.001; “****“, *P* < 0.0001. D Extracted ion chromatograms of terrestric acid (top), plipastatin B2 (middle) and surfactin C14 (bottom) detected in *P. hordei* and P5_B1 co-culture on LB maltose.

Here, the four main organic acids produced by *P. hordei* steadily increase over time, peaking on day 7 and 9 and reaching concentrations around 3-fold higher than in monocultures (TA on day 9, t-test, *P* = 0.0000532; CA on day 9, t-test, *P* = 0.00217) (Fig. 2C and Fig. S4).

We corroborated such observation by propagating both interacting partners on LB agar plates supplemented with 20% maltose, a growth condition where acid production is highly reduced, but the LP production is maintained to a level similar to the ones obtained on PDA plates (Fig. 2B and Fig. 2D). Under this condition, we demonstrated that lipopeptide precipitation via acid production confers a protective mechanism for *P. hordei.* In the absence of such precipitation, *P. hordei* growth was impaired by *B. subtilis*, as evidenced by the clear inhibition halo (Fig. 2B).

### *Corymbifera* series penicillia induce lipopeptide precipitation via different organic acids

To investigate how widespread the white line precipitation phenomenon is, we co-inoculated *B. subtilis* with eight fungal strains closely related to *P. hordei* from the *Corymbifera* series on pH indicator plates, as described above. To capture a broad range of fungal diversity, we also included distantly related fungal species from *Penicillium* series *Adametziorum, Viridicata, Atroveneta, Scabrosa, Soppiorum, Olsoniorum, Camembertiorum, Citrina* and *Westlingiorum* [7]. All the *Corymbifera* species tested, except *Penicillium hirsutum* , induced *B. subtilis* L P precipitation, mirroring the abundance and spatio-temporal precipitation patterns observed in the interaction with *P. hordei* (Fig. 3A). A targeted metabolomics approach, combined with known chemotaxonomic markers [7,35,36], revealed that fungi capable of inducing the white precipitate produced various organic acids, including TA, CA, and VA acid (Fig. 3A). Within the *Corymbifera* series, *Penicillium albocoremium* induced white line precipitation by predominantly producing barceloneic acid A, whose concentration was approximately 17-fold higher than viridicatic acid. Two other *Corymbifiera* species exhibited contrasting precipitation patterns: *P. hirsutum* did not induce LP precipitation, consistent with its low TA and CA acid production compared to *P. hordei* (one-way ANOVA, *P*lil=lil2.3lil×lil10⁻¹⁴) (Fig. 3B-D), whereas *Penicillium allii* triggered precipitation despite lacking TA and its biosynthetic derivates and producing only fulvic acid (Fig. 3A and Fig. S5).

**Fig. 3.**
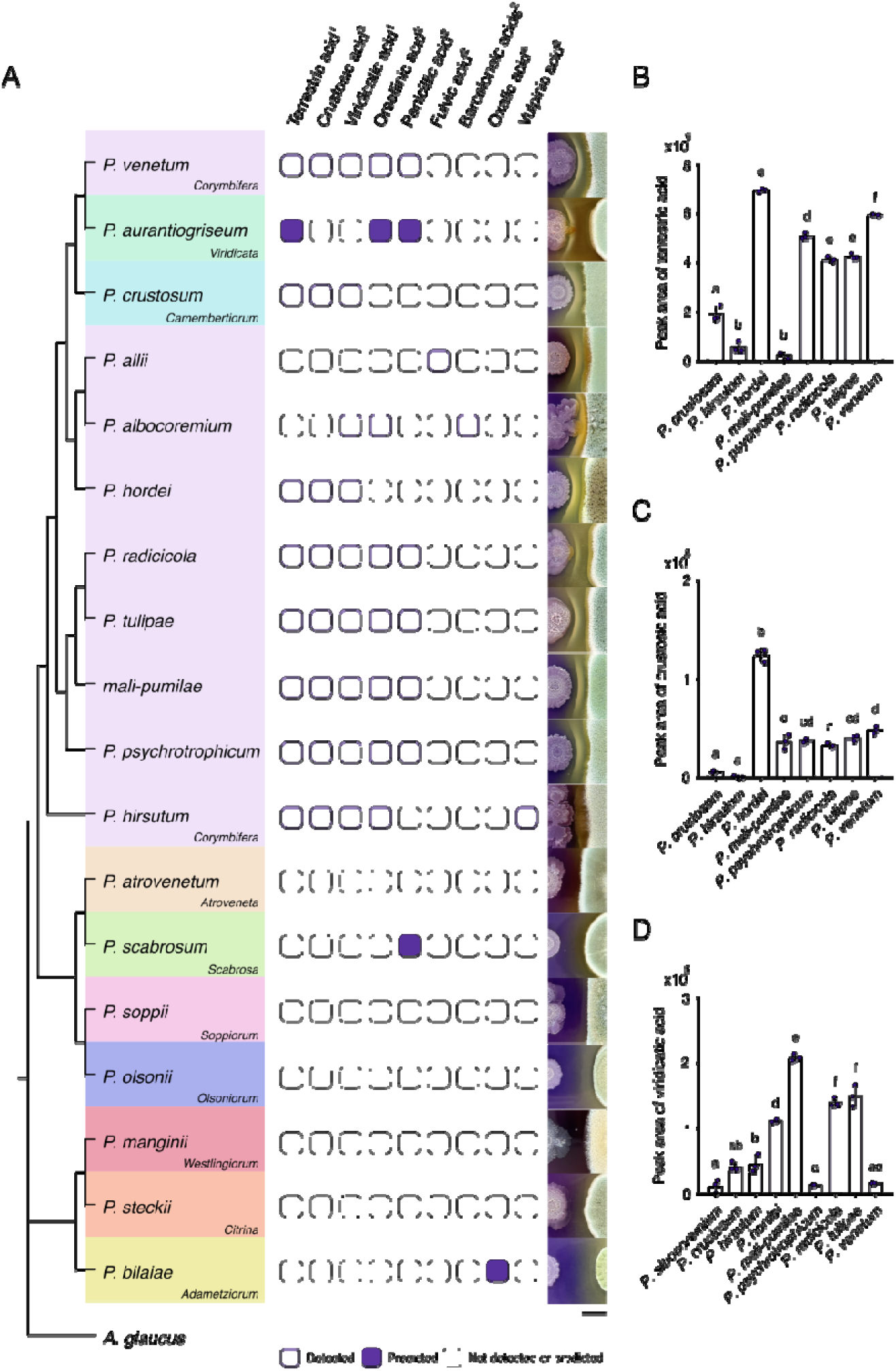
*Corymbifera* series penicillia induce lipopeptide precipitation through secretion of distinct organic acids. **A** Closely related *Corymbifera* species except for *P. hirsutum* induced lipopeptide precipitation via production of terrestric, fulvic or barceloneic acids. Among distantly related fungi, *P. crustosum and P. aurantiogriseum* were also able to form the white precipitate through the producing terrestric acid. The phylogenetic tree was built based on calmodulin (CaM) sequences of interacting type fungal strains using maximum likelihood (ML) method and Kimura (1980) 2-model [37]. The phylogram was rooted with *Aspergillus glaucus* as an outgroup and the analysis was conducted in MEGA12 [38]. Fungal species are color-coded according to their taxonomic series, indicated in the lower-right corner of each colored box. Light purple squares indicate acids detected by LC-MS, dark purple squares indicate acids predicted to be produced by certain fungal species based on chemotaxonomic survey, and white squares indicate that no acids were detected or predicted. Numbers next to acids refer to identification level (1-3): (1) reference standard, (2) confident match based on MS/MS and (3) MS only [39]. The letter a indicates compounds identified as chemotaxonomic markers based on a chemotaxonomic survey. Images on the right display the presence or absence of the white precipitate between *B. subtilis* P5_B1 and the respective fungal species on pH indicator plates. Scale bar = 5 mm. B-D Production of terrestric, crustosic and viridicatic acids of *Corymbifera* and *Camembertiorum* species varied in co-culture with *B. subtilis*. Acid production was quantified by LC-MS (peak area) and mean values were calculated from three biological replicates (n = 3). Error bars represent standard deviation; grouping letters are from ANOVA with Tukey–Kramer’s post hoc test, where different letters within plots indicate a statistically significant grouping on day 9 (P1<10.05).

Among distantly related species tested, only *Penicillium crustosum* (*Camembertiorum* series) and *Penicillium aurantiogriseum* (*Viridicata* series), both members of the *Fasciculata* section, were able to induce LP precipitation through the production of TA. Other distantly related *Penicillium atrovenetum, Penicillium scabrosum, Penicillium olsonii, Penicillium soppii, Penicillium manginii, Penicillium steckii* and *Penicillium bilaiae* were all able to modify the pH of the interaction plates yet failed to precipitate LPs. The chemotaxonomic survey showed that apart from *P. scabrosum* producing penicilic acid, none of these species are known to produce the organic acids detected in the *Corymbifera* species. Overall, these data confirm that fungal species capable of inducing white precipitation also produce higher levels of multiple organic acids, linking acid production to the observed white line phenomenon.

### Plipastatin production is reduced by terrestric acid-mediated acidification

Once we determined the direct role of *P. hordei* acidification in causing LP precipitation, we investigated its consequences for *B. subtilis,* given that lipopeptides are crucial defensive metabolites in bacilli. To do so, we confronted *P. hordei* with a panel of *B. subtilis* strains on pH indicator plates: the parental wild type (WT), *sfp* (impaired in non-ribosomal peptide synthesis), *srfAC* (surfactin-deficient), and the double mutant *srfAC* Δ*ppsC* (lacking both plipastatin and surfactin). In co-cultures with the WT strain, plipastatin production declined significantly, dropping to about 30% of monoculture levels by days 5 and 7 (t-test, *P* = 0.00045 and *P* = 0.00528) (Fig. 4A). When acidification was reduced on LB-maltose medium, plipastatin levels rose on day 9 (t-test, *P* = 0.03), and *B. subtilis* more effectively inhibited *P. hordei*, suggesting that fungal mediated acidification impacts plipastatin production (Fig. 4A).

**Fig. 4.**
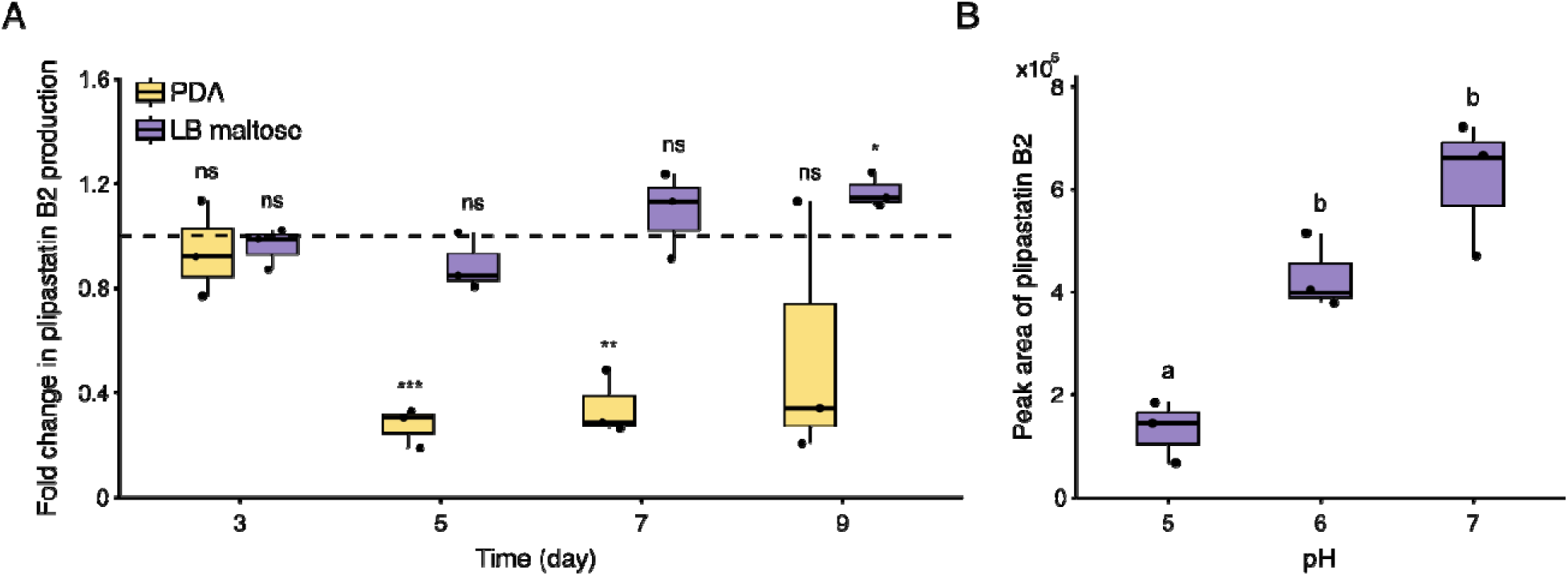
Plipastatin production is reduced by fungal-mediated acidification. **A** Impact on plipastatin B2 production in the presence of fungal acidification on PDA (yellow) and under highly reduced acidification on LB maltose (purple). Fold changes were calculated based on relative quantification of the plipastatin B2 by LC-MS (peak area) in co-culture compared to monoculture of *B. subtilis* P5_B1 (fold change = 1, dashed line). Box plots were generated from three biological replicates (n = 3), whiskers span from the minimum to maximum values, and black line inside the box indicates the median. Statistical significance was calculated using Student’s t-test where “ns” denotes no significant difference; "*“, *P* < 0.05; "**“, *P* < 0.01; "***“, *P* < 0.001. **B** Plipastatin B2 production by *B. subtilis* P5_B1 decreased when grown alone on PDA at low pH values (pH 5, 6 and 7) on day 7. Plipastatin B2 production was quantified by LC-MS (peak area) from three biological replicates (n = 3) per each pH value; grouping letters are from ANOVA with Tukey–Kramer’s post hoc test, where letters within a plot indicate a statistically significant grouping (*P*l1<l10.05).

To confirm that acidification leads to reduced plipastatin production, we cultivated the *B. subtilis* WT strain alone on plates with different starting pH values (pH 5, 6 and 7). On day 5, plipastatin levels were comparable across all pH conditions (one-way ANOVA resulted in nonsignificant differences between groups) (Fig. S6). In contrast, on day 7, plipastatin levels at pH 5 were significantly reduced than those at pH 6 and 7 (one-way ANOVA, Plil=lil1.8lil×lil10⁻⁴), whereas no significant difference was observed between pH 6 and 7 (Fig. 4B).

We next asked whether the reduction in plipastatin production was a general response to low pH, regardless of which acid-producing fungal species *B. subtilis* was confronted with. Closely related fungi and *P. crustosum* were co-cultivated for 9 days in darkness as described above. Here, plipastatin production was strongly reduced during interactions with TA-producing species, including *P. venetum* , *P. tulipae* , *P. radicicola* , and *P. psychrotrophicum (t-test, P = 0.035, P = 0.0053, P = 0.013, P = 0.017)* (Fig. S7). However, enhanced plipastatin production was observed upon co-cultivation with *P. allii* and *P. albocoremium* , both species that lack TA production but instead produce fulvic and barceloneic acids, respectively (Fig. 3A and Fig. S5). TA and its associated intermediates show the ability to affect plipastatin production, whereas other organic acids fail to do so despite their induced lipopeptide precipitation, indicating that *Bacillus* exhibits a species-dependent response to acidic conditions.

### Surfactin-mediated precipitate acts as a protective barrier

We then examined the contribution of surfactin to the white precipitate formation and its respective role in the interaction. In contrast to plipastatin, fungal acidification had no significant impact on surfactin production in WT strain (Fig. S8). When *P. hordei* was confronted with *srfAC* strain, a white line precipitate formed near fungal edge, but it was less intense compared to that formed in the interaction with the WT strain (Fig. 5A). As expected, MALDI-MSI analysis revealed accumulation of only plipastatins *(m/z* 1477.8288, 1491.8231, and 1505.7993 were identified as proton adducts of plipastatin B2, plipastatin A2 and plipastatin B1) at the white precipitate region. These ions showed the highest signal at the white precipitate region, with a reduced signal on the outer side of the *Bacillus* colony, facing away from the fungus (Fig. 5B). The loss of surfactin production resulted in stronger inhibition of the fungus compared to the WT strain, as indicated by denser and elevated aerial hyphae and the accumulation of brown exudate droplets at the interface (Fig. 5A), which formation is often associated with cell death or stress response. Compared to the WT strain, *srfAC* strain was more affected and exhibited an increased swarming motility between day 7 and day 12. Altogether, these observations confirm the vital role of surfactins in a white precipitate formation, serving as a physical barrier that provides protection for both interacting partners.

**Fig. 5.**
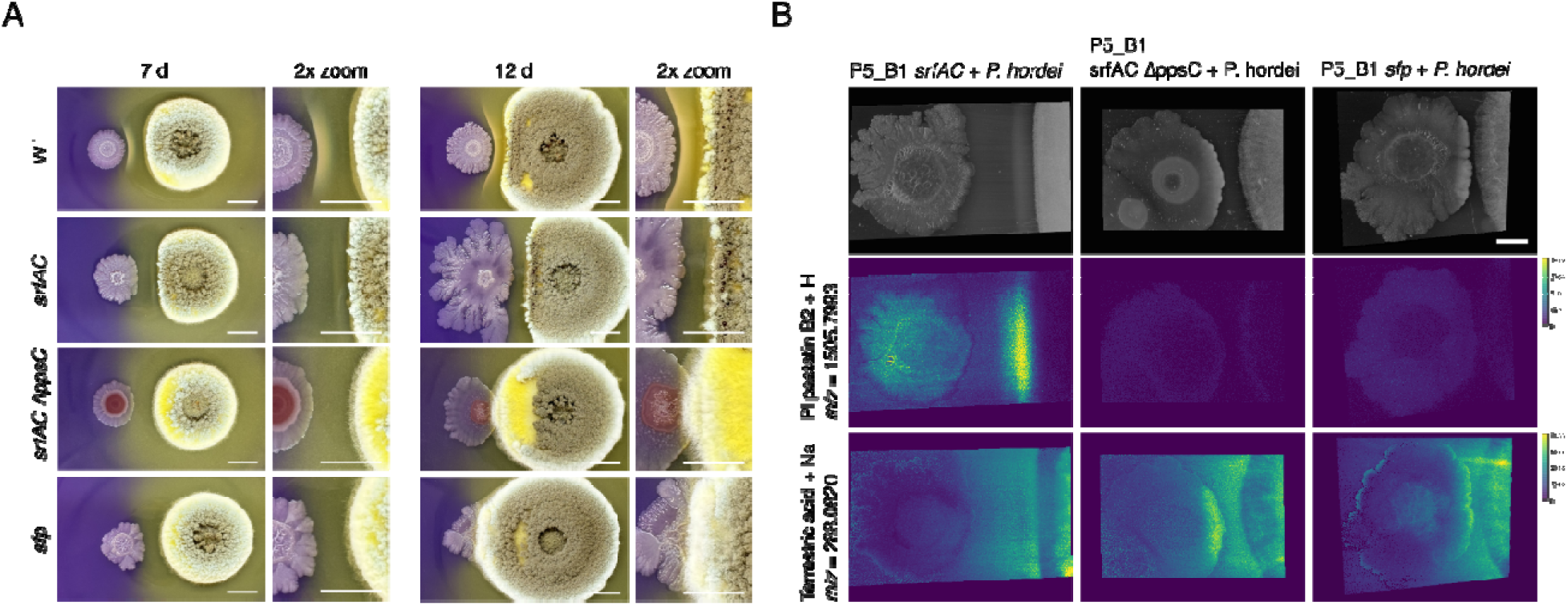
Lack of lipopeptide production by *B. subtilis* allows *P. hordei* to invade and overgrow *the Bacillus* colony. **A** Time-course of interactions between *P. hordei* and *B. subtilis* P5_B1 WT, P5_B1 *srfAC*, P5_B1 *srfAC* Δ *ppsC* and P5_B1 *sfp* on day 7 and day 12 on pH indicator plates. Close-up (2x) of interacting area is shown on the right. Left colonies are *B. subtilis* strains and right colonies are *P. hordei*. Scale bar = 1 cm. **B** MALDI-MSI reveals the spatial distribution of plipastatin B2 and terrestric acid in the interactions with *B. subtilis* mutants on day 9. Ion images are shown in viridis color scheme and colored scale bars besides ion images indicate total ion count. Scale bar = 5 mm (top-right image).

### B. subtilis relies on lipopeptide production for defense

Since the *srfAC* strain was still able to resist fungal takeover, we co-cultivated *P. hordei* with *srfAC* Δ*ppsC* and *sfp* strains. In both cases, *P. hordei* was able to partially invade the frontline of *Bacillus* colony after 12 days and after 15 days, nearly the entire colony was covered with *P. hordei* mycelium (Fig. 5A, Video S2). Compared to interactions with the WT and *srfAC* strains, the distribution pattern of TA (*m/z* 233.0820 [M+Na]^+^) was altered with increased signal intensity at the frontline of the *Bacillus* colony, suggesting localized accumulation (Fig. 5B). This was further supported by the yellow coloration of the outer edge of *Bacillus* colony at the interface. Despite lacking lipopeptide production, the mutant strains retained the ability to enhance TA production in the fungus. Both the *srfAC* Δ*ppsC* and *sfp* strains increased TA levels to a similar extent as the WT strain on day 9, thereby ruling out lipopeptides as the inducing signals (Fig. S9). Interestingly, without the lipopeptides, *B. subtilis* lacks the ability to buffer the medium, which allows for TA to diffuse through the bacterial colony. Subsequently, to evaluate if TA is toxic for *B. subtilis*, we tested five different concentrations of pure TA (400, 200, 100, 50, and 25 µg mL ^-1^) on *B. subtilis* strains (WT, *srfAC* and *sfp*) and their growth was monitored over 24 h period. The results showed that TA at all tested concentrations had no inhibitory effect on *B. subtilis* growth (Fig. S10).

Having established that white line precipitation is required for *B. subtilis* to prevent its overgrowth, we next examined whether *P. hordei* could cause the same effect when confronted with other LP-producing bacteria. While *P. hordei* was able to precipitate lipopeptides in co-culture with *Bacillus velezensis* and *Bacillus tequilaensis* , it failed to do so with all tested *Pseudomonas* strains (Fig. S11). In contrast to bacilli, *Pseudomonas* strains were able to acidify the culture medium and did not exhibit any deleterious effects on fungal growth.

## Discussion

In this study, we investigated interaction between *B. subtilis* , known for its strong microbial antagonistic activity, and the non-pathogenic, rhizosphere-associated *P. hordei*. Cross-kingdom interactions between fungi and bacteria are crucial for community functionality and actively shape ecological niches, yet SMs driving these interactions remain underexplored [6,40]. We demonstrate for the first time that *Penicillium*–*Bacillus* interaction leads to precipitation of lipopeptides (plipastatins, surfactins and gageopeptides), forming insoluble aggregates, observed as a white line precipitate in agar. Although the white line phenomenon has been reported and is presumed to result from the co-precipitation of CLPs between interacting *Pseudomonas* strains, the underlying molecular mechanism remains unclear [41–44].

Our findings show that co-cultivation with *B. subtilis* led *P. hordei* to enhance its production of fungal organic acids, particularly TA and its biosynthetic derivatives. When acid production was highly reduced through medium composition modifications, *P. hordei* failed to precipitate secreted lipopeptides, indicating that fungal organic acids drive the white line precipitation phenomenon. Mechanistically, bacterial lipopeptide precipitation likely follows principles like those used in methods for recovering lipopeptides from cell-free broths via acidification. In this process acid neutralizes the negative charges on lipopeptides, reducing their solubility in the aqueous phase and inducing precipitation [45,46]. In the absence of the precipitated lipopeptides, *B. subtilis* was able to inhibit the growth of *P. hordei*, suggesting that the presence of the acids serves as vital protective mechanism for the fungus. We propose that fungal organic acids induce lipopeptide precipitation to mediate fungal defense in two distinct ways: chemical inhibition and physical barrier formation.

*B. subtilis* produces plipastatins known for their strong antifungal properties against a wide range of fungi [47,48]. Plipastatin, like fengycin and surfactin, acts on fungal membranes, causing structural damage and altering their permeability, which can result in cytoplasm leakage and ultimately lead to cell death [49–51]. Amino acid sequences of the cyclic core and variations in fatty acid chain lengths have a significant impact on the antifungal activity of lipopeptides, where even slight structural changes can affect their activity [52]. As a result, the insoluble lipopeptide aggregates upon acidification therefore cannot reach the fungal colony to exert their antifungal activity.

We observed that accumulation of lipopeptides over time results in a pronounced white precipitate that forms a physical barrier, clearly separating *Penicillium* from *Bacillus*. Although in the interaction with surfactin production-impaired mutant ( *srfAC*) white precipitate, composed of plipastatins, was formed, it was not as strong as the one in the co-culture with wild type. Plipastatin-mediated precipitate failed to form a distinct barrier, which resulted in stronger fungal inhibition. The observed variation in precipitate formation could be attributed to differences in production levels between plipastatin and surfactin. Since surfactin is produced in much higher amounts than plipastatin, more surfactin molecules are available to precipitate, which leads to a more pronounced white precipitate [53]. This suggests that surfactins are potential key contributor to the white line precipitate which protects the fungus by limiting the diffusion of antifungal plipastatins and other metabolites toward the fungal colony.

In contrast to plipastatin, surfactin is not a directly antifungal metabolite in biologically relevant concentrations [54], however, it plays a crucial role in *Bacillus* fitness in the rhizosphere by promoting motility, biofilm formation, and root colonization [21,55,56]. Hong-Wei and Ming Chu reported that surfactin starts to precipitate out of the production medium at pH lower than 5 and redissolves completely when the pH returns to 6.1 [57]. Due to its smaller size and shorter fatty acid tail, we assume surfactin is more prone to precipitation than plipastatin and acts as a starting point for the formation of the white precipitate barrier. Interestingly, the loss of surfactin production was associated with increased *B. subtilis* sensitivity to *P. hordei* , suggesting that surfactin-mediated barrier indirectly shields *Bacillus* from *Penicillium* S M s . Similarly, Andrić et al. showed that white precipitate formed through LP-aggregation in the interaction between *Pseudomonas* and *Bacillus* serves as a protective mechanism to shield *Bacillus* from sessilin-mediated toxicity [58]. Therefore, we ascertain that precipitation of surfactin provides a competitive advantage to *P. hordei* by impairing *Bacillus* motility and root colonization to enable the fungus to secure its niche and nutrients in the rhizosphere.

Surfactin and plipastatin are viewed as two key metabolites produced by *Bacillus* to thrive in competitive microbial interactions [59]. *P. hordei* was able to invade and eventually overgrow *Bacillus* mutants impaired in either surfactin and plipastatin production ( *srfAC* Δ*ppsC*) or in the production of surfactin, plipastatin, bacillaene and bacillibactin ( *sfp*). MSI analysis revealed accumulation of TA at the frontline of the *B. subtilis* mutant colony, especially in *srfAC* Δ*ppsC* mutant, indicating that TA may promote the fungal penetration and overgrowth of the bacterial colony. TA was found to be cardiotoxic and displays concentration-dependent phytotoxic effects on rice seedlings [60,61]. Although TA showed no toxicity to *B. subtilis* at tested concentrations, it is possible that higher concentrations accumulated in the interaction could exert inhibitory effect on the *Bacillus* growth.

Beyond lipopeptide precipitation, fungal acidification significantly reduced plipastatin production. Reduction of bioactive metabolite production via acidification has been demonstrated and was suggested to provide a general advantage against microbes producing antibiotics which are less active at acidic pH [62]. The decrease in plipastatin production was a TA-mediated response, whereas acidification via other organic acids led to enhanced plipastatin production. This suggests that *B. subtilis* responds to acidification in a fungus-specific manner and that low pH is not the sole factor affecting the production of bioactive SMs. In line with our results, previous studies showed that production of antifungal LPs varies among *Bacillus* species and is highly influenced by the presence of fungal competitors [63–65]. *Bacillus* can either overproduce antifungal lipopeptides to outcompete the interacting fungi or their production is suppressed in response to specific signals emitted by fungi, thereby potentially promoting fungal-bacterial co-existence.

Lipopeptide precipitation was conserved among *Corymbifera* species through production of TA, fulvic or barceloneic acids, indicating a restricted phylogenetic distribution of such defense strategy and a limited set of organic acids capable of causing this effect. However, non-*Corymbifera* species *P. crustosum* and *P. aurantiogriseum*, both of which produce TA, were also able to induce white line precipitation. This suggests that TA at certain concentrations may act as a universal metabolite driving LP precipitation, thereby pointing to a broader distribution of such defensive strategy among penicillia to counteract bacilli and persist in the niche to exert potential beneficial effects on the plant.

Our findings emphasize the importance of environmental cues and fungal signals in regulating SM production in *Bacillus* and offer insights into the broader concept of acidification as a microbial defense strategy.

## Data availability

FBMN workflows can be found at the GNPS website under the following links:

(1) *P. hordei*-*B. subtilis* co-culture on PDA (positive mode): https://gnps.ucsd.edu/ProteoSAFe/status.jsp?task=8e0f80d153d34f3b894f4eb36d3724ca;
(2) *P. hordei*-*B. subtilis* co-culture on PDA (negative mode): https://gnps.ucsd.edu/ProteoSAFe/status.jsp?task=c32e6695bddf4b86aa7f3f6d8707a82c;
(3) *Corymbifera*-*B. subtilis* interactions (positive mode): https://gnps.ucsd.edu/ProteoSAFe/status.jsp?task=67d33af1d1c0471ea1cb943a5be62099;
(4) *Corymbifera*-*B. subtilis* interactions (negative mode): https://gnps.ucsd.edu/ProteoSAFe/status.jsp?task=7eab8907eb45410b9b4df511907ef765
(5) *P. hordei*-*B. subtilis* co-culture on LB maltose (positive mode): https://gnps.ucsd.edu/ProteoSAFe/status.jsp?task=02592e5e63344fe5a10d7463b559c777

## Author contributions

Conceptualization: MV, JCF, CNLA, and SAJ; Experimentation: MV, MDS, AJCA; Writing: MV, CNLA, and SAJ, with edits from all co-authors.

## Supporting information

Timelapse of wild types P. hordei and B. subtilis

Timelapse of wild type P. hordei and B. subtilis srfAC

## Acknowledgements

MV, AJCA, and JCF were supported by the Novo Nordisk Foundation (NNF19SA0059360). MDS, CNLA and SAJ were supported by the Danish National Research Foundation (DNRF137) for funding the Center for Microbial Secondary metabolites (CeMiSt). We would also like to acknowledge funding from Novo Nordisk Foundation (grant NNF19OC0055625) for the infrastructure “Imaging microbial language in biocontrol (IMLiB) and to CNLA via the grant NNF24OC0093315. Finally, we acknowledge the DTU Metabolomics Core for usage of LCMS equipment as well as daily maintenance.

