## Supplementary Data for "Penicillium hordei acidification precipitates Bacillus subtilis lipopeptides to evade inhibition"

**SUPPLEMENTARY INFORMATION**

***Bacillus subtilis* lipopeptide precipitation induced by *Penicillium hordei* acidification prevents fungal inhibition**

Manca Vertot^a^, Morten D. Schostag ^a^, Aaron J. C. Andersen^a^, Jens C. Frisvad^a#^, Carlos N. Lozano-Andrade ^a#^, Scott A. Jarmusch ^a#^

^a^Department of Biotechnology and Biomedicine, Technical University of Denmark, Lyngby, Denmark

Includes Supplementary Figures S1-S11, Supplementary Tables S1-S2, Supplementary Videos S1-S2, and Supplementary References

**SUPPLEMENTARY FIGURES**

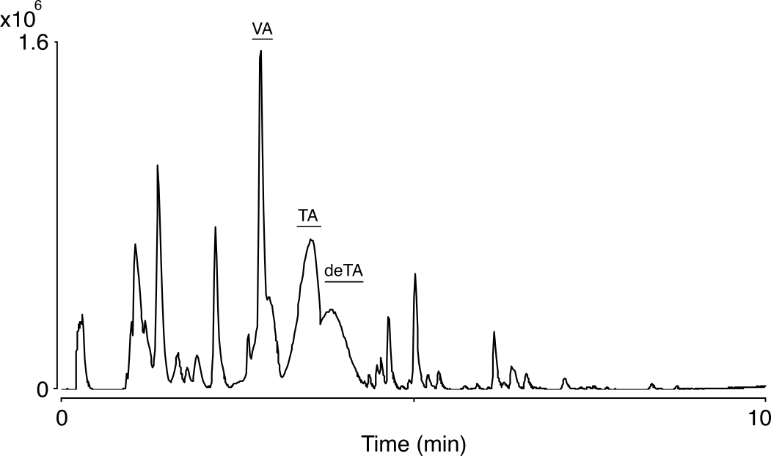

**Fig. S1.** **LC-MS of the white line precipitate.** Base peak chromatogram (BPC) in negative mode of the crude extract from the white precipitate region area showing the accumulation of fungal organic acids. Letters above peaks correspond to the following acids: VA: viridicatic acid, TA: terrestric acid, deTA: dehydroterrestric acid. The y-axis represents peak intensities.

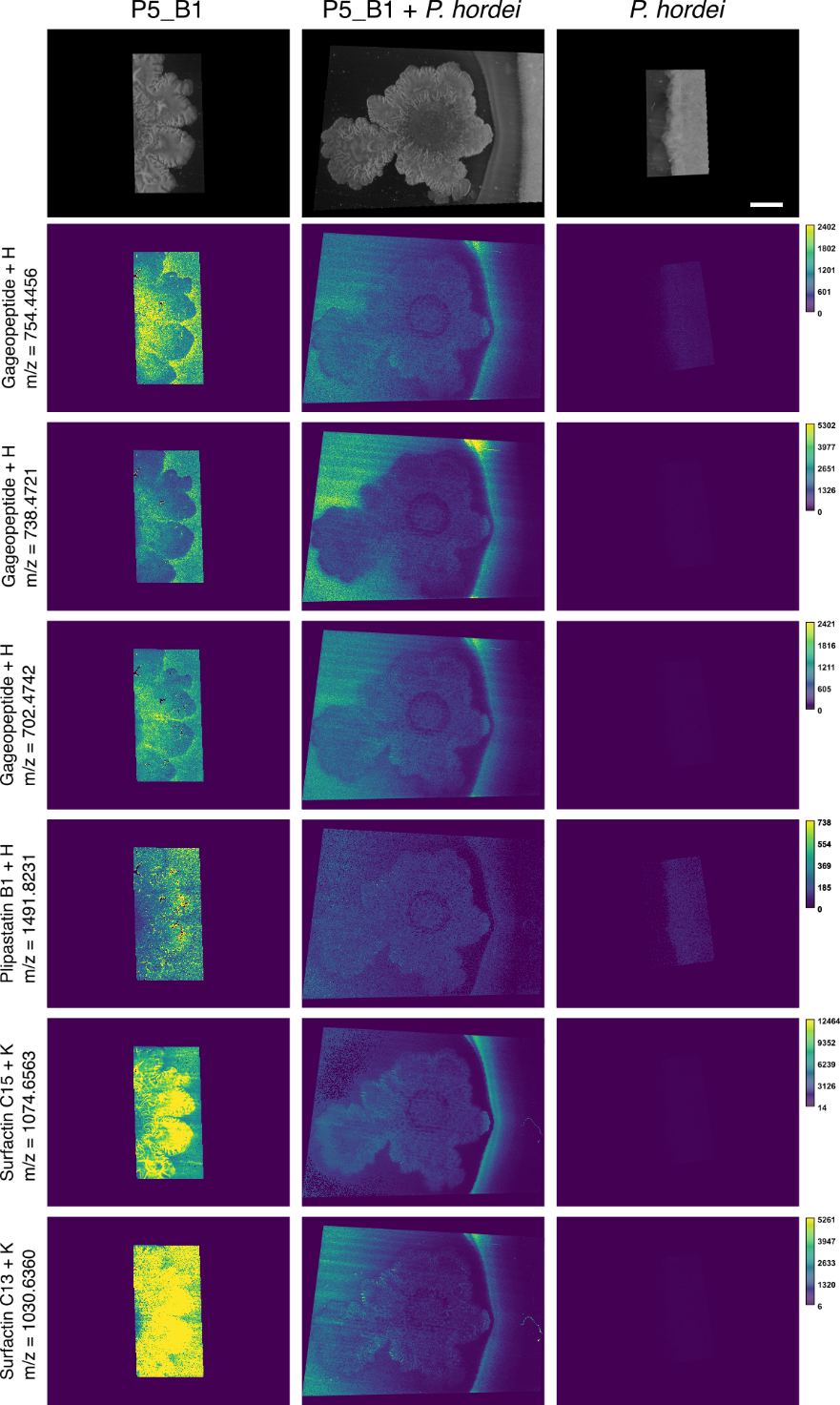

**Fig. S2.** **MALDI-MSI of selected ions in the interaction and corresponding monocultures.** Ion images are shown in viridis color scheme and colored scale bars besides ion images indicate total ion count. Scale bar = 5 mm (top-right image).

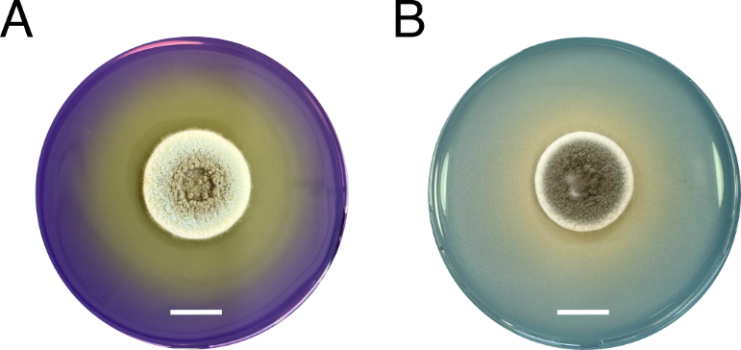

**Fig. S3. Extracellular acidification ability of *P. hordei.* A** *P. hordei* grown alone on PDA supplemented with bromocresol purple for 7 days. A purple color indicates a pH above 6.8, while a yellow color indicates a pH below 5.2. **B** *P. hordei* grown on PDA supplemented with bromocresol green for 7 days. A green color indicates a pH above 5.2, while a yellow color indicates a pH below 3.8. Scale bar = 1 cm.

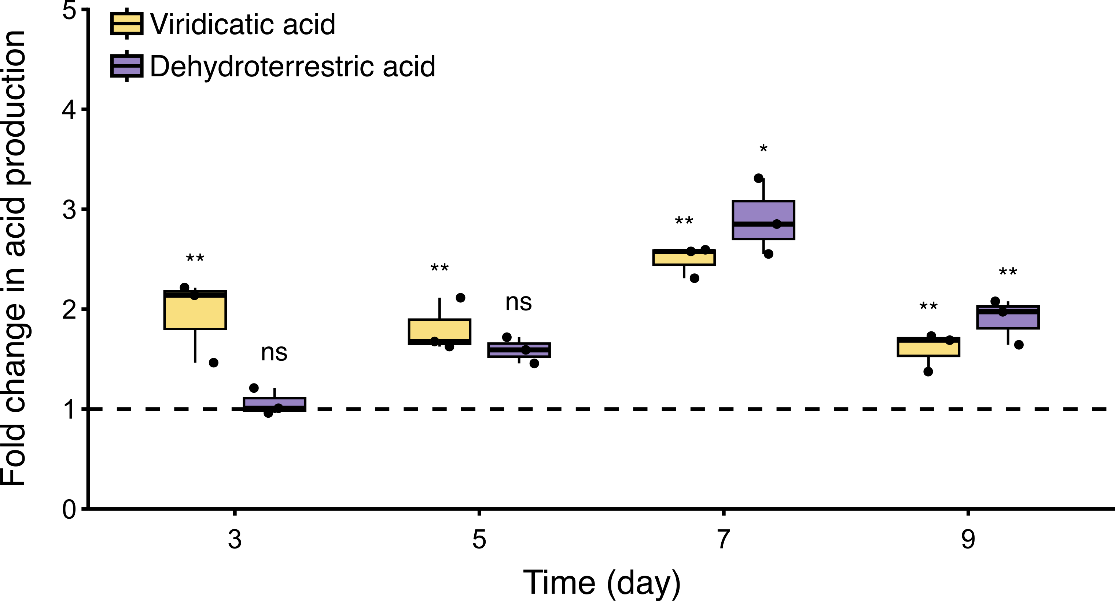

**Fig. S4. Impact on acid production by *P. hordei* in the interaction with *B. subtilis.*** Viridicatic (yellow) and dehydroterrestric (purple) acid production by *P. hordei* in response to *B. subtilis* over time (days 3, 5, 7, 9) on pH indicator plates. Fold changes were calculated based on relative quantification of the acids by LC-MS (peak area) in co-culture compared to monoculture of *P. hordei* (fold change = 1, dashed line). Box plots were generated from three biological replicates (n = 3), where whiskers span from the minimum to maximum values, and black line inside the box indicates median. Statistical significance was calculated using Student’s t-test where “ns” denotes no significant difference; "*“, *P* < 0.05; "**“, *P* < 0.01.

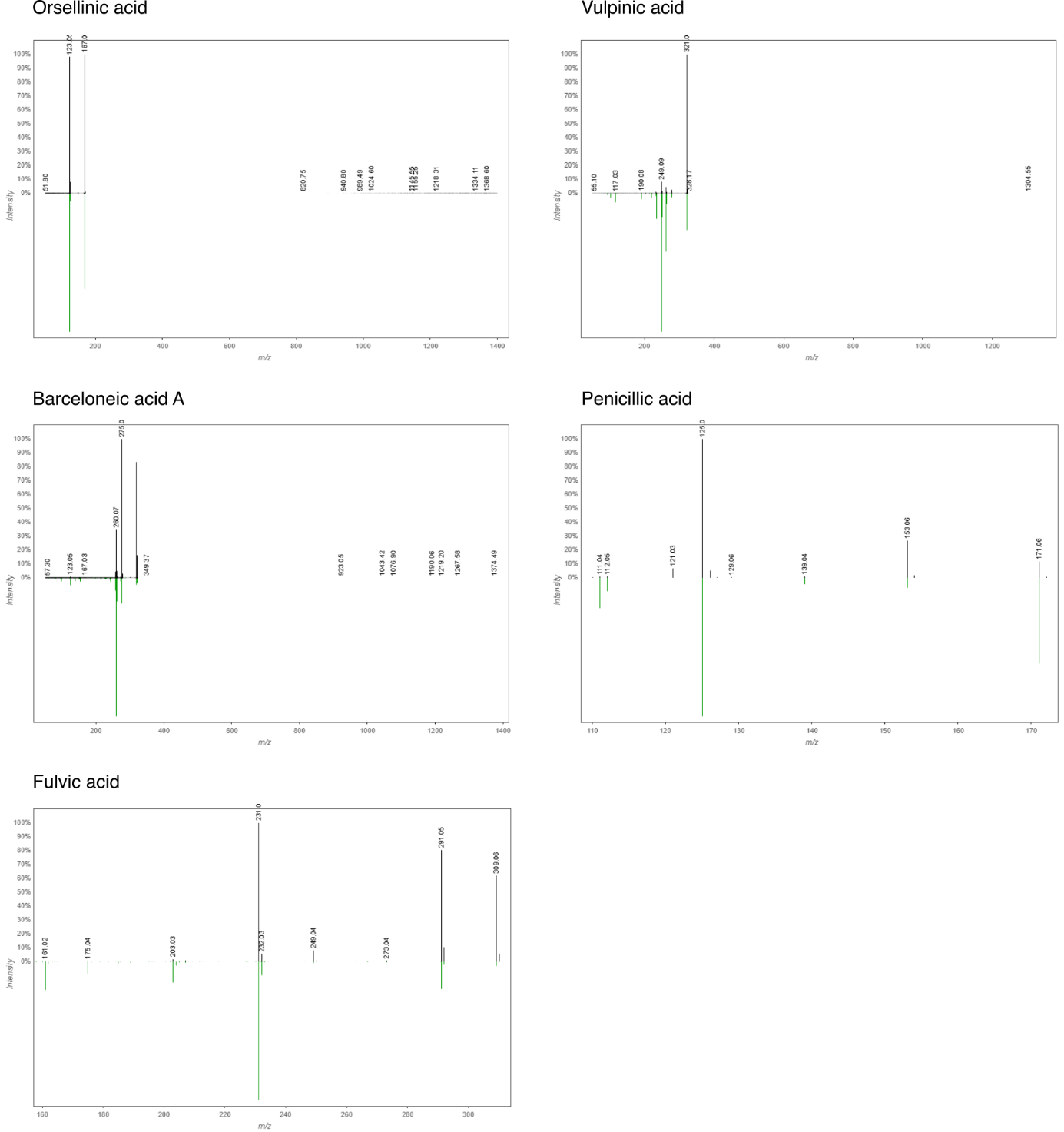

**Fig. S5. Mirror plots showing fragmentation library matches for detected acids.** The green (bottom) spectrum represents the library reference, while the black (top) spectrum corresponds to the experimental data.

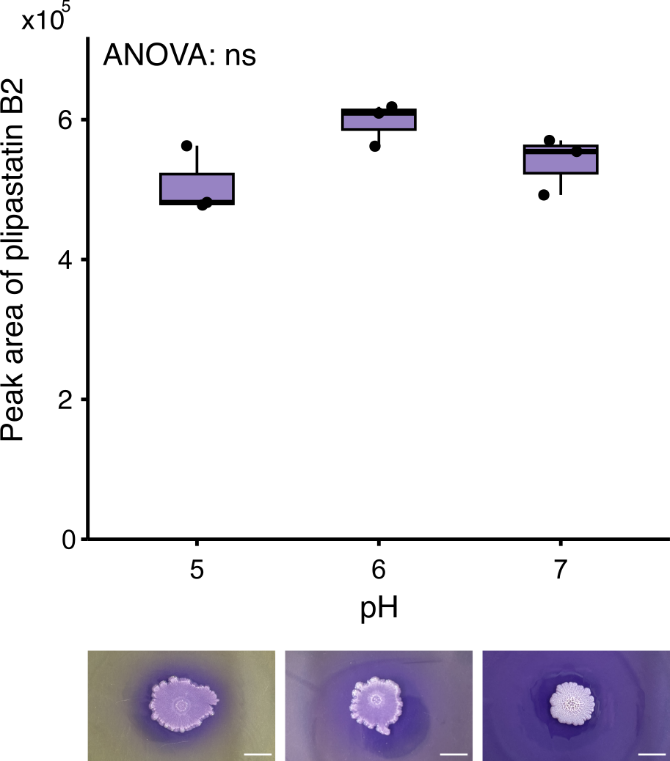

**Fig. S6. Plipastatin B2 production by *B. subtilis* at different pH values on day 5***. B. subtillis* was cultivated on PDA at pH values (pH 5, 6 and 7) for 5 days. Images of *B. subtilis* P5_B1 on pH 5 (left), pH 6 (middle) and pH 7 (right); scale bar = 1cm. Plipastatin B2 production was quantified by LC-MS (peak area) from three biological replicates (n = 3) per each pH value.

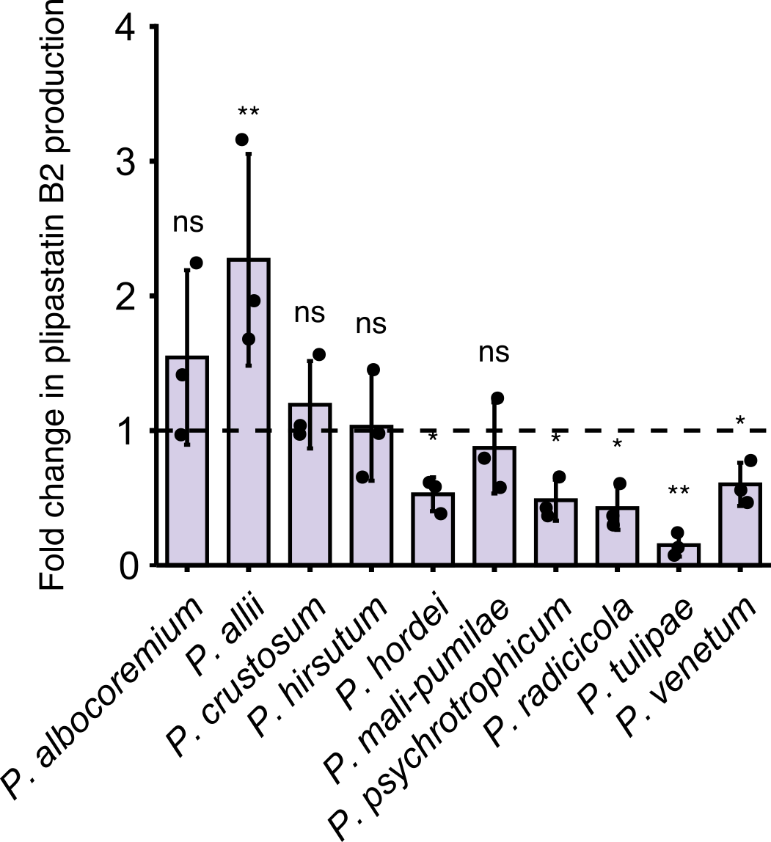

**Fig. S7. Changes in plipastatin B2 production by *B. subtillis* in interaction with *Corymbifera* and *Camembertiorum* species.** Fold changes were calculated based on relative quantification of the plipastatin B2 by LC-MS (peak area) in co-culture compared to monoculture of *B. subtilis* P5_B1 (fold change = 1, dashed line) on day 9. Mean values were calculated from three biological replicates (n = 3) and error bars represent standard deviation (n = 3). Statistical significance was calculated using Student’s t test where “ns” denotes no significant difference; "*“, *P* < 0.05; "**“, *P* < 0.01.

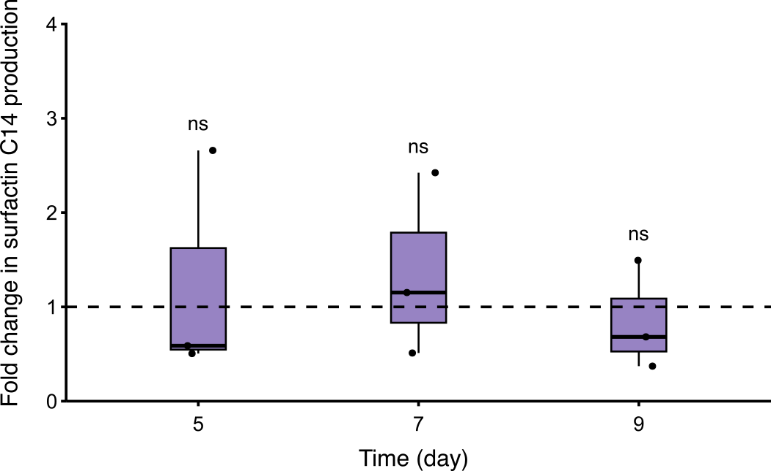

**Fig. S8. Impact on surfactin production by *B. subtilis* in the interaction with *P. hordei*.** Surfactin C14 production by *B. subtilis* in response to *P. hordei* over time (days 3, 5, 7, 9) on pH indicator plates. Fold changes were calculated based on relative quantification of a surfactin C14 by LC-MS (peak area) in co-culture compared to monoculture of *B. subtilis* P5_B1 (fold change = 1, dashed line). Box plots were generated from three biological replicates (n = 3), whiskers span from the minimum to maximum values, and black line inside the box indicates median. Statistical significance was calculated using Student’s t-test where “ns” denotes no significant difference.

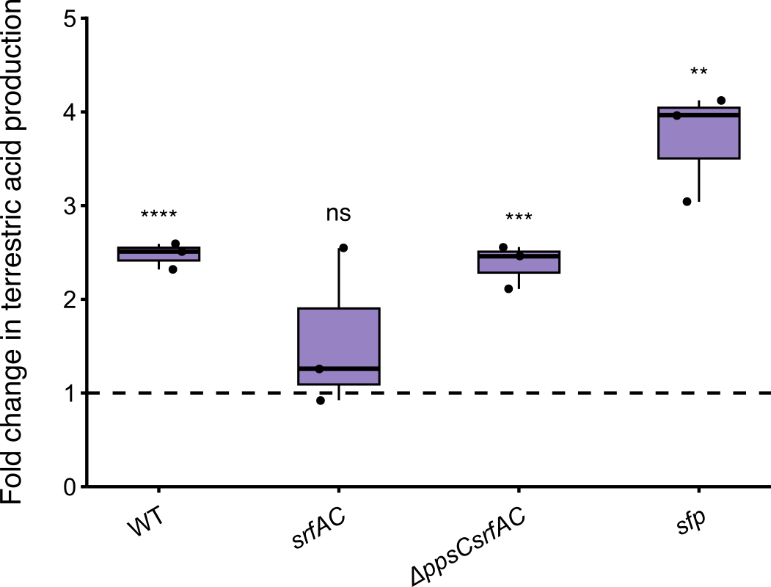

**Fig. S9. Impact on terrestric acid production by *P. hordei* in response to *B. subtilis* mutants*.*** *P. hordei* was co-cultivated for 9 days with the following *B. subtilis* strains: the parental wild type (WT), *sfp* (impaired in non-ribosomal peptide synthesis), *srfAC* (surfactin-deficient) and the double mutant *ΔppsC srfAC* (lacking both plipastatin and surfactin). Fold changes were calculated based on relative quantification of terrestric acid by LC-MS (peak area) in co-culture compared to monoculture of *P. hordei* (fold change = 1, dashed line). Box plots were generated from three biological replicates (n = 3), where whiskers span from the minimum to maximum values, and black line inside the box indicates median. Statistical significance was calculated using Student’s t-test, where “ns” denotes no significant difference; "**“, *P* < 0.01; "***“, *P* < 0.001; “****“, *P* < 0.0001.

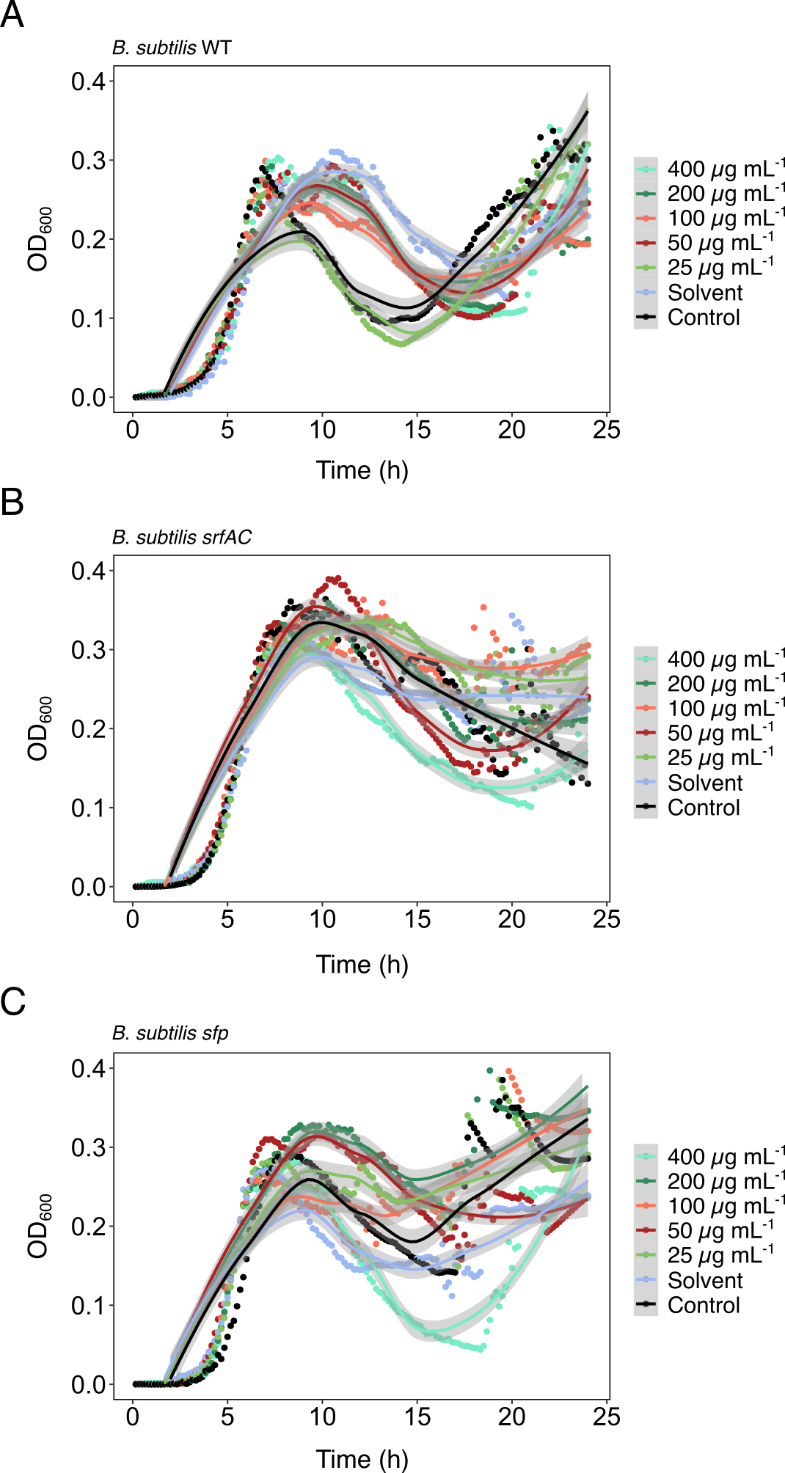

**Fig. S10. Effect of terrestric acid on the growth of *B. subtilis* strains.** *B. subtilis* (**A** *B. subtilis* WT, **B** *B. subtilis* *srfAC* and **C** *B. subtilis* *sfp*) was grown in LB media supplemented with varying concentrations of terrestric acid (25 to 400 µg mL^-1^), dissolved in 50:50 H_2_O:MeOH solution. LB served as a control, whereas LB supplemented with methanol served as the solvent control (Solvent). The growth was measured over 24 hours using OD_600_ and the first measurement (Time = 0) was subtracted from all following measurements. The hard lines represent an estimated growth curve, whereas the dots are the average of four replicates (n = 4). One replicate of *B. subtilis srfAC* in LB supplemented with 400 µg mL^-1^ terrestric acid was removed due to contamination.

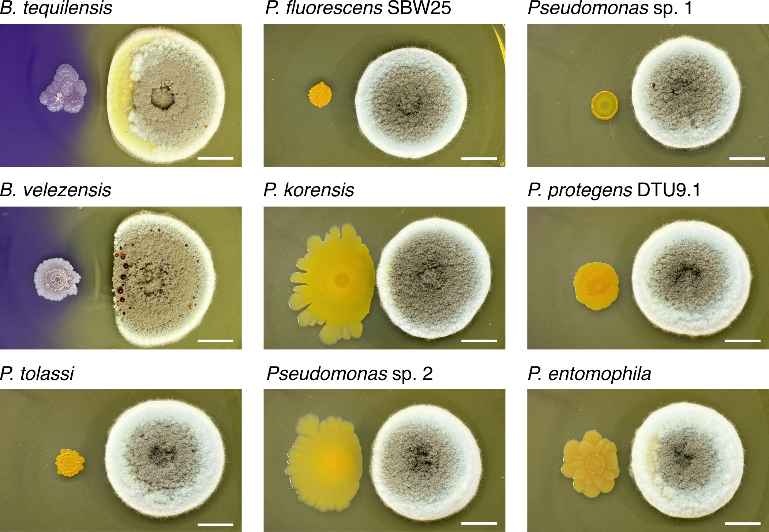

**Fig. S11. Interaction between *P. hordei* and other lipopeptide-producing bacteria.** Bacterial strains were co-cultured with *P. hordei* on pH indicator plates for 9 days. Scale bar = 1 cm.

**SUPPLEMENTARY TABLES**

**Table S1.** Bacterial strains used in this study.

| **Strains** | **Genotype** | **Ref.** |
| --- | --- | --- |
| ***Bacillus subtilis*** |  |  |
| P5_B1 | Wild type | [1] |
| P5_B1 *srfaAC* ::Tn10 (Spec^R^) | P5_B1 disrupted *srfAC* gene; unable to produce surfactins | [1] |
| P5_B1 ∆*sfp*::mls | *Sfp* mutant; unable to produce NRPs and NRPs/PKs | [1] |
| P5_B1 *srfaAC* ::Tn10 (Spec^R^); ∆*ppsC* (Tet^R^) | P5_B1 disrupted *srfAC* gene and deleted *ppsC* gene; unable to produce surfactins and plipastatins | This study |
| 3610 ∆*ppsC* (Tet^R^ ) | 3610 deleted *ppsC* gene; unable to produce plipastatins | [1] |
| ***Bacillus velezensis*** | Wild type | In-house collection |
| ***Bacillus tequilensis*** | Wild type | In-house collection |
| ***Pseudomonas protegens*** DTU9.1 | Wild type | [2] |
| ***Pseudomonas entomophila*** | Wild type | In-house collection |
| ***Pseudomonas korensis*** | Wild type | In-house collection |
| ***Pseudomonas* sp. 1** | Wild type | In-house collection |
| ***Pseudomonas* sp.** **2** | Wild type | In-house collection |
| ***Pseudomonas fluorescens*** SBW25 | Wild type | In-house collection |

**Table S2**. Fungal strains used in this study.

| **Strains** | **IBT number** |
| --- | --- |
| ***P. hordei*** | IBT 35745 |
| ***P. crustosum*** | IBT 36642 |
| ***P. allii*** | IBT 20212 |
| ***P. albocoremium*** | IBT 22806 |
| ***P. venetum*** | IBT 10591 |
| ***P. tulipae*** | IBT 22526 |
| ***P. psychrotrophicum*** | IBT 33673 |
| ***P. radicicola*** | IBT 10696 |
| ***P. mali-pumilae*** | IBT 33672 |
| ***P. hirsutum*** | IBT 10623 |
| ***P. manginii*** | IBT 36724 |
| ***P. atrovenetum*** | IBT 36306 |
| ***P. scabrosum*** | IBT 36325 |
| ***P. olsonii*** | IBT 35623 |
| ***P. soppii*** | IBT 36723 |
| ***P. steckii*** | IBT 36728 |
| ***P. bilaiae*** | IBT 13516 |
| ***P. aurantiogriseum*** | IBT 35768 |
